# Co-adaptation of dynamic human-machine interfaces

**DOI:** 10.1101/2023.07.14.549053

**Authors:** Amber H.Y. Chou, Momona Yamagami, Maneeshika M. Madduri, Benjamin J. Chasnov, Finley Hutchison, Lauren N. Peterson, Samuel A. Burden

## Abstract

Despite the growing prevalence of learning algorithms in daily life, methods for analysis and synthesis of how these systems interact with people are limited. We studied optimization-based algorithms that co-adapt with people in the presence of dynamic machines, finding limitations on current theory that motivated us to conduct an experiment where human subjects interact through a dynamic interface with a machine that has complex dynamics. Experimental results provided evidence of co-adaptation and a trade-off between performance and the “effort” of the human and interface, defined as the norm of their output signals. We developed a parsimonious model of the human adaptation strategy observed in our experiments and conducted a simulation study using this model. Our computational results matched the empirical results, suggesting our human subjects adapted to minimize a combination of error and effort. These results demonstrate how co-adaptation between humans and intelligent interfaces shapes behavior and performance, and introduces a modeling framework that can be used in future work to systematically design interaction outcomes.

## I. Introduction

With practice and effort, people can learn to control dynamic systems like vehicles [1]–[3], tele- and co-robots [4]–[6], prostheses [7]–[9], exoskeletons [10]–[12], or brain-computer interfaces [13]–[15]. To facilitate and shape this learning, it is tempting to inject intelligence into the *interface* between the human and machine [16]. Since people continually adapt to their sensorimotor context [17], [18], an intelligent interface in this closed-loop interaction creates a two learner problem [19], [20], where the human and interface *co-adapt* [21]. Analyzing and synthesizing these systems is critically important in current and emerging applications, including driver assistance [22], surgical robotics [23], rehabilitative robotics [24], active prosthetics [8], and neural interfaces (both invasive [13] and non-invasive [25], [26]).

Motivated by the optimal feedback control hypothesis for human motor control [27]–[30], we modeled the interaction between a human, adaptive interface, and dynamic machine using a robust control framework [31]. We assumed that both the human and the interface solve optimal control problems to determine their behaviors. In our theoretical analysis, we allowed arbitrarily large (but finite) system orders of the dynamic machine, and analyzed the equilibrium behavior of such systems. We found that a simple form of this problem has no solution due to a fundamental theoretical limitation neglected in prior work [32]–[34]. In prior work, it was assumed that the human and the interface had noise-free observations of the machine’s state vector, which we regard as unrealistic in real-world applications. For example, in the context of neuroprosthetics, where an intelligent neural interface seeks to help a person control a dynamic device, intrinsic stochasticity of neural signals [35] injects noise into the system, precluding access to full information. This finding led us to conduct a human subjects experiment and simulation to understand and model how people behave in-the-loop with adaptive interfaces and dynamic machines under more realistic conditions.

In our experiment, we investigated co-adaptation between humans and adaptive interfaces in the presence of specific machine dynamics that are fundamentally challenging to control [2]. We recruited participants (*N* = 11) to perform a continuous disturbance-rejection task using a one-dimensional manual input device [3]. We deployed adaptive interfaces with 0th-, 1st-, and 2nd-order dynamics, which correspond to the position-, velocity-, and acceleration-based systems that people routinely encounter in daily life [1]. The interface controller coefficients were updated as the participant completed a sequence of trials by estimating a model of the participant’s feedback controller and optimizing a performance criterion of interest. We compared the system performance and human behavior with co-adaptation and without co-adaptation, finding that co-adaptation led to diverse outcomes exhibiting trade-offs between agent behavior and system performance.

Driven by this experimental finding, we conducted a computational analysis to test whether a model for co-adaptation can replicate the empirical outcomes we observed. Specifically, we modeled human behavior as arising from minimization of a cost function. We implemented a simulation where the human model and interface alternately optimize their costs and we conducted a parameter sweep over the space of penalty terms in the two agents’ cost functions. This simulation yielded behaviors that aligned closely with empirical observations. Our experimental and computational findings will inform future frameworks and deployments of co-adaptive interfaces in human-machine interactions.

## II. A model of co-adaptation

We consider systems where the machine has dynamics, that is, where the dimension of the machine’s state vector – the system *order* [36, Sec. 2.2] – is greater than or equal to 1. The zero-dimensional case was studied in prior work [37]– [39]. We modeled dynamic human-machine interfaces (HMI) as the block diagrams in Figure 1, where: *H* represents the *human* “in-the-loop”, *M* is the *machine* that is being controlled, and *I* is the *interface* we seek to synthesize. When the blocks *H, M*, and *I* are given linear time-invariant (LTI) transformations [40, Lec. 3], these diagrams are not solely conceptual – they provide mathematical specifications of the closed-loop transformation from input disturbance *w* to output error *z*. Although humans are generally nonlinear, they can behave remarkably linearly when interacting with finite-order LTI machine-and-interface dynamics [1]. These observations motivate us in what follows to focus on the finite-order LTI case where there exists a comprehensive toolkit for analysis and synthesis of feedback systems [31], [36], [40], [41].

**Fig. 1.**
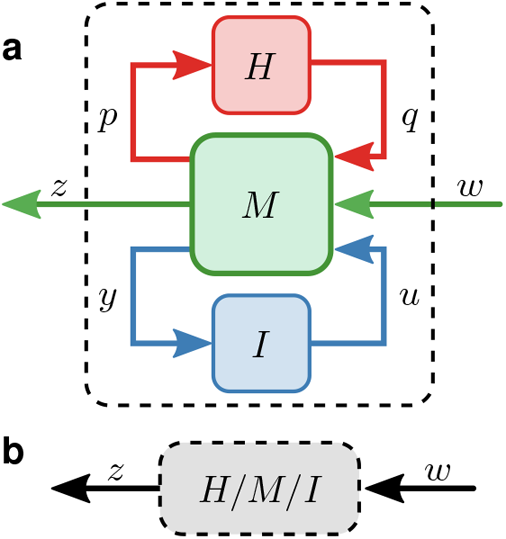
Block diagram models for human-machine interfaces (HMI). (**a**) This diagram specifies that the human *H* transforms signal *p* to signal *q*, i.e. the human observes output *p* from machine *M* and provides control input *q* to the machine. Similarly, the interface *I* transforms machine output *y* to control input *u*, and the machine *M* transforms both of the control inputs *q, u* and an external input *w* to produce outputs *p, z, y*. The external input *w* can contain a *disturbance* to reject (e.g. measurement or process noise) or a *reference* to track (e.g. a trajectory or stationary point). (**b**) This diagram shows a simplification of (a) that obscures details about the interconnection between *H, M*, and *I*, instead denoting the transformation *H/M/I* resulting from this interconnection. The block *H/M/I* is defined so that (a,b) both specify the same transformation from *w* to *z*.

### A. Interface synthesis

The field of *robust control theory* [31] provides methods for synthesizing the interface *I* in Figure 1 (conventionally termed the *controller*) to optimize a performance criterion with respect to fixed models of human *H* and machine *M* (and, optionally, uncertainty in the models). The performance criteria of interest in robust control are system norms that quantify how much signal power is transferred from input disturbance *w* to output error *z*. In this robust control paradigm, the interface *I* is synthesized by solving an optimization problem

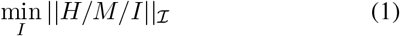

where the norm ∥·∥_ℐ_ encodes what components of *z* the interface seeks to make as small as possible. Under appropriate restrictions on the choice of norm ∥·∥_ℐ_, models of human *H* and machine *M*, and statistics of input *w* and output *z*, a solution *I*^∗^ to the optimization problem in (1) exists [31] and can be computed using efficient numerical algorithms [41]. As an example, the well-known *linear-quadratic Gaussian* (LQG) regulator [40, Lec. 24] is obtained by solving (1) when: the disturbance *w* is Gaussian; the error *z* consists of (linear transformations of) the state of the machine *x* and the control input *u*; the human-machine transformation *H/M* (defined in Figure 2) is stabilizable and observable [40, Lec. 14, 15]; and ∥·∥_ℐ_ is the 2-norm.

**Fig. 2.**
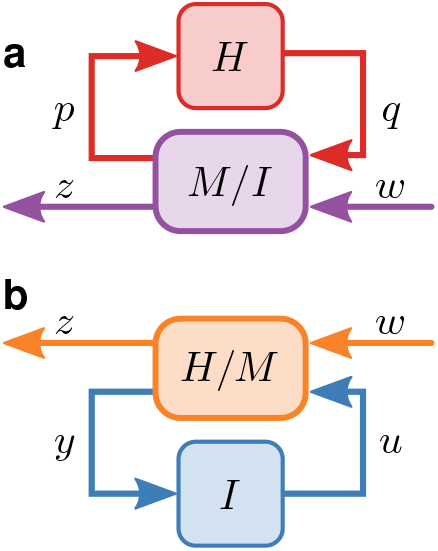
Block diagrams from human and machine perspectives. Given LTI transformations *H, M*, and *I*, the blocks *M/I* and *H/M* are defined so that the diagrams in (**a, b**) specify the same transformation from *w* to *z* as the diagrams in Fig. 1. Mathematically, the *U/L* operation is defined as the *linear fractional transformation* [31, Ch. 3, 10] between blocks *U* and *L*. These simplified diagrams are conceptually useful when reasoning from the individual perspectives of the human *H* and interface *I* as they jointly interact with the machine *M*. Indeed, *H* interacts with the interconnection *M/I* between *M* and *I* as illustrated in (a), whereas *I* interacts with the interconnection *H/M* between *H* and *M* as in (b).

If the statistics of the human *H* and machine *M* in Figure 1 are stationary regardless of the implemented interface *I* (e.g. if *H* and *M* are given as fixed LTI transformations or distributions of LTI transformations), then the preceding paragraph describes a remarkably flexible framework for synthesizing an optimal interface *I*^∗^ [31], [42]. However, although it may be reasonable to assume or ensure machine dynamics are stationary, ample evidence suggests that the human will naturally adapt to any perceived change in the interface, and moreover that this adaptation will not be random – rather, the human’s transformation will be strongly influenced by the interface [1], [2], [25], [43]. Therefore if the interface *I*^∗^ is synthesized by solving (1) with respect to an initial guess or estimate of the human’s transformation *H*, it is reasonable to expect the human will adapt its transformation from *H* to 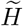 when the interface changes from *I* to *I*^∗^. Unfortunately, the synthesized interface *I*^∗^ is not optimal with respect to the adapted human transformation 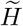. In fact, implementing 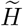 in-the-loop with *H* could yield arbitrarily bad performance [44].

### B. Co-adaptation

The preceding observations motivate the study of interfaces that *co-adapt* with the human and, hence, regard the human *H* and interface *I* as two *learners* [19], [20] playing a *dynamic game* [45] through their interaction with the machine *M*. As a starting point for modeling this interaction, prior work suggests the human may play this game by solving their own optimization problem [27]–[30]

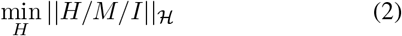

where the norm ∥·∥_ℋ_ encodes what components of *z* the human seeks to make as small as possible. Other than in the special case where the goals of the human and interface are perfectly aligned so that ∥·∥_ℐ_= ∥·∥_ℋ_, the outcome of the game defined by simultaneously considering the optimization problems in (1) and (2) will generally represent a compromise between the player’s conflicting goals. One such outcome considered in prior work on human-machine interfaces [19], [32], [33], [37] is a *Nash equilibrium* [45], [46] defined by a pair of transformations *H*^∗^, *I*^∗^ such that *H*^∗^ minimizes ∥*H/M/I*^∗^∥_ℋ_ with respect to *H* and *I*^∗^ minimizes ∥*H*^∗^*/M/I*∥_ℐ_ with respect to *I*.

### C. Non-existence of Nash equilibria

This section provides theoretical analysis of the dynamic game defined in the preceding section that is played by a human *H* and interface *I* interacting with a machine *M*. We will assume in what follows that *H, M*, and *I* are finite-order linear time-invariant (LTI) transformations of continuous-time signals. The game of interest is specified by the coupled optimization problems

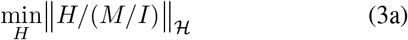

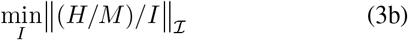

where ∥·∥_ℋ_, ∥·∥_ℐ_ denote the cost functions of the human and interface, respectively, and the closed-loop transformations *H/*(*M/I*) = *H/M/I* = (*H/M*)*/I* are defined in Fig. 2. The simplified block diagrams in Fig. 2 are useful to illustrate (i) the combined machine-interface system (*M/I*) the human *H* interacts with and (ii) the combined human-machine system (*H/M*) the interface *I* interacts with.

Given LTI transformations *M* and *I*, the (central, strictly feasible) solution *H*^∗^ of the optimization problem in (3a) with respect to the 2-or ∞-norm is, under standard stabilizability and detectability conditions, an LTI transformation [31, Thm 14.7] that can be computed using efficient algorithms [41]. With the exception of the special cases considered in prior work [32]–[34] (termed *full information* and *full control* in [31], [47]), the number of state variables in the dynamics of *H*^∗^ (the system’s *order*) is equal to the sum of the number of state variables in *M* and *I*; if we let #(*T*) denote the number of state variables in the LTI transformation *T*, this property can be written #(*H*) = #(*M*) + #(*I*). Similarly, given LTI transformations *H* and *M*, the solution *I*^∗^ of the optimization problem in (3b) is an LTI transformation, and the number of state variables in *I*^∗^ equals the sum of the number of state variables in *H* and *M* : #(*I*) = #(*H*) + #(*M*). We will assume this generic condition holds in what follows.

We will show in the Theorem below that the game in (3) generally has no Nash equilibrium defined in terms of finite-order LTI transformations when *M* has dynamics, i.e., #(*M*) ≥ 1. To see why this may be the case, consider the co-adaptive interaction wherein *H* and *I* alternately solve their optimization problems ((3a) and (3b), respectively). Even starting from a static interface with #(*I*) = 0, solving (3a) with respect to given *M* and *I* yields a solution *H*^∗^ with #(*H*) = #(*M*) ≥1, so subsequently solving (3b) with respect to given *H*^∗^ and *M* yields solution *I*^∗^ with #(*I*) = #(*H*) + #(*M*) = 2 #(*M*) ≥ 2. Iterating this process yields a sequence of *H* and *I* with ever-increasing numbers of state variables, preventing the existence of a stationary point for (3) in the sense of Nash defined with finite-order LTI transforms.

#### Theorem

Suppose M is a finite-order linear time-invariant (LTI) transformation with dynamics so that 1 ≤ #(M) < ∞, and suppose that ∥·∥_ℋ_ and ∥·∥_ℐ_ are system 2-or ∞ -norms. If there exists a Nash equilibrium for (3) defined in terms of finite-order LTI transformations H^∗^, I^∗^, then at least one of H^∗^/M or M/I^∗^ is full information or full control.

#### Proof (by contradiction)

Suppose there exists a Nash equilibrium *H*^∗^, *I*^∗^ for (3) such that *H*^∗^ and *I*^∗^ are LTI and have finite order. If neither *H*^∗^*/M* nor *M/I*^∗^ is full information or full control, then [31, Thm. 14.7] implies #(*H*^∗^) = #(*M*) + #(*I*^∗^) and #(*I*^∗^) = #(*H*^∗^) + #(*M*). Substituting the second equation from the preceding sentence into the first and simplifying yields 0 = 2 #(*M*). But since #(*M*) ≥ 1, this equation is a contradiction.

#### Remark

The co-adaptation games studied in prior work [19], [32]–[34], [37] consider only the full information case, where it is assumed that the interface can observe the state of the machine M and human H with no measurement noise and similarly the human can observe the state of the machine and interface I with no noise. In this case, the game can admit a Nash equilibrium defined in terms of static state feedback transformations for H and I [45, Sec. 6.2.2].

#### Example

The neuroprosthetic example from the introduction illustrates why full information or full control assumptions are unrealistic. Indeed, full information would require both adaptive agents – human and interface – have noise-free measurements of all system states, including those internal to the other adaptive agent. Similarly, full control for either agent would require that agent to directly influence all system states, including those internal to the other; although it might be possible in principle to give the brain full control over the machine and interface, it seems implausible (and perhaps undesirable) for the interface to have full control over the human’s neural state. Either assumption seems inconsistent with our understanding of neural dynamics [35].

#### Remark

The preceding Theorem only applies to the case where H, M, and I are all finite-order LTI transformations as in [31]. Indeed, solutions of multi-agent stochastic optimal control problems need not be linear even if dynamics are linear and disturbances are Gaussian [48]; we cannot say at present whether the human-machine game considered here has nonlinear stationary points. Furthermore, the techniques in [31] are not applicable in the presence of delay or other common structural constraints, which necessitate the use of new tools [42].

## III. Methods

The experiment and computational analysis were designed to determine whether and how human-machine interfaces (HMI) co-adapt, and to characterize how co-adaptation affects human, interface, and system performance relative to a non-adaptive *baseline*.

### A. Human subjects

Eleven participants were recruited (age: 19-36; gender: 6 women, 5 men; hand dominance: 10 right-handed, 1 left-handed). All were daily computer users. All participants provided informed consent according to the University of Washington, Seattle’s Institutional Review Board (IRB #00000909).

### B. Task

Participants controlled a one-dimensional (1D) cursor using a 1D slider (35× 12 ×22 mm rectangular handle on a 10 cm slide potentiometer; Figure 3a). The slider input *u*_*H*_ was transformed through interface *I* and a fixed machine *M* to produce cursor position *y* on the screen (Figure 3c). Participants were instructed to keep the cursor as close to the center line, *y* = 0, as possible (Figure 3b). We chose a nonminimum-phase second-order machine to increase task complexity [2]:

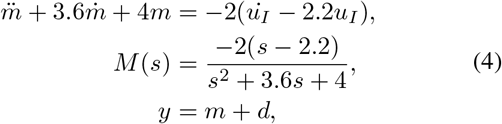

where 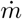 represents the time-domain machine output *m* differentiated by time, *M* (*s*) is the continuous-time representation of transformation *M* in the frequency domain, *s* = *jω*. The cursor position *y* was updated at 60 Hz, so the continuous-time machine was discretized (scipy.signal.cont2discrete).

**Fig. 3.**
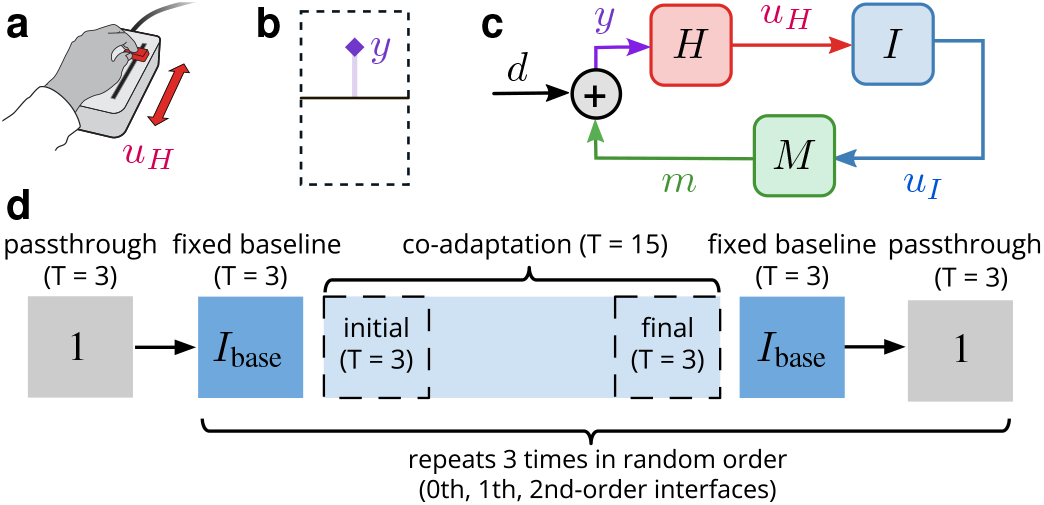
Human subjects experiment design. (**a**) Participants provide output *u*_*H*_ using a 1D manual device to (**b**) control the position of a cursor *y* on a computer display to minimize the distance to a reference line in the middle of the screen. (**c**) Human *H*, machine *M*, and interface *I* are connected in series, with the human viewing cursor *y* and producing response *u*_*H*_ that is input to *I*. The machine output is perturbed by external disturbance signals *d*. (**d**) Experiment conditions and trial structure. *T* indicates the number of trials (e.g., *T* = 3 means there are 3 trials in the block). Three conditions (0th-, 1st-, and 2nd-order interface) are presented in random order, followed by 3 pass-through trials after each block.

We introduced an unpredictable disturbance *d* to the cursor (Figure 3c), which mimics the measurement or process noise from external devices. Following prior work [3], [26], [49], disturbance signals were constructed as a sum of sinusoidal signals with the first eight prime multiples of a base frequency of 1*/*20 Hz (*stimulated frequencies* Ω = [0.10, 0.15, 0.25, 0.35, 0.55, 0.65, 0.85, 0.95] Hz). The magnitudes at each frequency component was normalized by its frequency to ensure constant signal power. The phase of each frequency component was randomized in each trial to produce pseudorandom time-domain signals.

### C. Signal processing

Prior work demonstrated that when humans are tasked with tracking references and rejecting additive disturbances through a linear time-invariant (LTI) [36, Ch. 3, pg. 4] system *M*, humans behave approximately like LTI transformations for a range of reference and disturbance signals [1], [3], [49]. Thus, we can analyze our system in Figure 3c using the frequency-domain representations [50, Ch. 5] of signals and LTI systems. In this paper, time-domain signal *x* with a “hat” 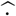 denotes the Fourier transform, 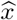, in the frequency-domain.

We recorded time-domain signals *y* and *u*_*H*_ at each sampling interval, and collected the updated interface dynamics *I* for each trial. In post-hoc analysis, we converted these signals into their frequency-domain forms, 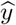 and 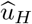, for each trial, then derived the interface input 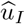 and the human transformation *H* (feedback controller) at stimulated frequencies:

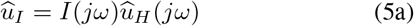

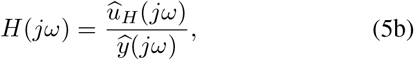

where *ω* ∈ Ω. Given this *H*, the prescribed signals *d*, and transformations *M, I*, we can additionally derive the steady-state signals via block diagram algebra [36, Sec. 2.2]:

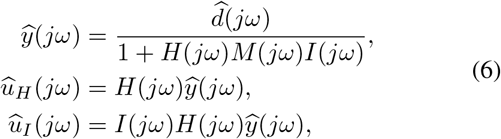

which were used in the interface updates (section III-E).

### D. Conditions

We tested interfaces that have 0th-, 1st- and 2nd-order dynamics (Table I), corresponding to the position-, velocity-, and acceleration-based systems [1] (Figure 3d). Participants first completed 3 trials of a pass-through interface with an identity mapping (*I*(*z*) = 1, i.e., *u*_*I*_ [*t*] = *u*_*H*_ [*t*]), then performed the three interface conditions in random order. Each condition block consists of 3 baseline trials, followed by 15 trials of co-adaptation, and 3 additional baseline trials. At the end of each condition block, participants completed another 3 pass-through trials, with a total of 12 pass-through trials per participant. Each trial was 40 seconds after a 5-second ramp-up (45 seconds total). Participants were encouraged to rest between trials and required to take a three-minute break between condition blocks. Total experimental duration was approximately 2 hours per participant.

### E. Interface adaptation

#### Interface cost

We defined an interface cost that minimizes human effort in addition to its own effort and task error [34]. The cost function was a linear combination of task error, interface effort, and human effort:

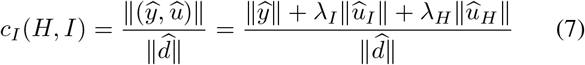

with penalty terms *λ*_*I*_ = 0.5 and *λ*_*H*_ = 1.5, which were estimated from pilot data to ensure the scaled interface and human efforts were within the same order of magnitude as the task error. We further scaled the interface penalty *λ*_*I*_ by half to avoid optimization from getting stuck at the smallest possible interface magnitudes. We used 2-norm to match the task instruction of minimizing the cumulative cursor displacement.

**TABLE I.**
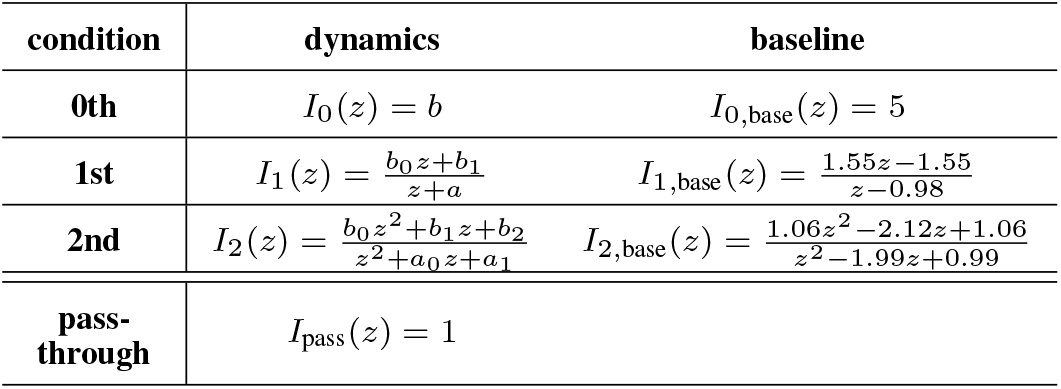
The discrete-time representations of 0th-, 1st-, 2nd-order, and pass-through interfaces tested in the experiment.z = e^jωdt^, where dt = 1/60 second.

#### Interface parameters discretization

The parameter space for each interface order was discretized into three candidate lists *I*_list_ of transfer-function coefficients (*a, b* in Table I).

##### 1) 0th-order interfaces

gain *b* ∈ [0.05, 5] to ensure interface remained responsive to user input without becoming unmanageably volatile, sampled at 100 equidistant points.

##### 2) 1st-order interfaces

We generated 1st-order high-pass interfaces by sweeping gains (0.05 to 5) and cutoff frequencies (0.05 to 1 Hz), with 100 equidistant points per range. The cutoff range was selected to align with the low-frequency crossover observed in human manual control tasks [3], [51].

Since the cursor updated at 60 Hz, we mapped continuous-time interface dynamics to their discrete-time representations. To ensure interfaces are stable or marginally stable, we excluded any interfaces where the eigenvalues of the denominator exceeded |1| in discrete time. We also excluded configurations resulting in negative DC gain at *z* = 1, to prevent control inversion, which our pilot participants reported as confusing and counter-intuitive. This resulted in a global parameter set of 8,546 combinations of coefficients (*b*_0_, *b*_1_, *a*).

##### 3) 2nd-order interface

We generated 2nd-order high-pass interfaces by sweeping gains (0.05 to 5), cutoff frequencies (0.05 to 1 Hz), and damping ratios (0.5 to 1), with 50 points per parameter for computational efficiency. Damping ratios below 1 were selected to ensure the interfaces remained stable and avoided aggressive noise amplification. We again computed the discrete-time representation of each interfaces, and then excluded configurations where interfaces were unstable or had negative DC gain, yielding a global set of 87,038 combinations of coefficients (*b*_0_, *b*_1_, *b*_2_, *a*_0_, *a*_1_).

For both 1st- and 2nd-order interfaces, we also explored low- and band-pass filters in pilot testing, but they consistently yielded higher task errors than high-pass filters. Thus, we restricted our adaptation framework to high-pass interfaces, as their phase-lead characteristic [52, Ch. 7.11] has the potential to compensate for sensorimotor delays in the nervous system.

#### Interface baselines

We started and ended each condition block with a baseline interface: 0th-order *I*_0,base_, 1st-order *I*_1,base_, and 2nd-order *I*_2,base_. Having different orders of baselines allowed for a controlled comparison within the same interface complexity, so we could verify whether improvements stem from co-adaptation or were simply an artifact of transitioning to a higher-order interface. We also use the baselines as the starting interfaces in co-adaptation.

We synthesized the baseline interfaces using pilot subject data, where 6 pilot subjects performed the same dynamic game with a pass-through interface (*I* = 1) and the second-order machine defined in (4). We estimated human transformations as (5b), then determined the optimal interface by searching through the parameter space *I*_list_ to minimize the interface cost *c*_*I*_ (*H, I*) in (7), for each of the three conditions. We used these three optimal interfaces as the baselines, and all participants used the same baselines.

#### Interface updates

The interface update occurred at the end of every co-adaptation trial for a total of 15 trials for each condition. The interface updates occurred in three steps.

##### 1) Human model

we estimated human transformation *H* in (5b) for each trial. This model was subsequently updated as a weighted combination between the historical estimate *H*^−^ and the current trial’s estimate *H*:

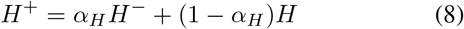

where *α*_*H*_ = 0.75 for gradual updates in the human model. For the first co-adaptation trial, the human model was initialized using data from the 3 preceding non-adaptive baseline trials.

##### 2) Interface grid search

For each candidate interface in *I*_list_, we computed the steady-state signals 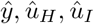 using (6) and generated *d* with random phase, then performed a grid search to identify a parameter set that minimizes the interface cost:

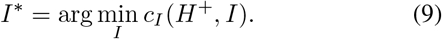

##### 3) Interface updates

*Smooth Batch* [13] was implemented to ensure gradual interface changes between trials. The interface for the subsequent trial *I*^+^ was defined as a weighted combination of the previous trial’s interface *I*^−^ and the optimal interface *I*^∗^:

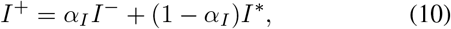

where *α*_*I*_ is the adaptation rate. We employed a conservative adaptation rate of *α*_*I*_ = 0.75 in this study. We selected this adaptation rate because previous studies using the Smooth Batch algorithm suggested that aggressive adaptation rates may lead to poor convergence [53]. A pseudo-code summarizing the steps of the interface update is in **Algorithm 1**.

#### F. Evaluation

We evaluated our experimental results against the following three hypotheses.

##### H1 (whether co-adaptation occurred)

We tested whether humans and interfaces changed after 15 trials of adaptation compared to non-adaptive baselines. For each condition (0th-, 1st-, 2nd-order), we compared the distributions (*N* = 11) of transformations, human ∥*H*∥ and interface ∥*I*∥, between the co-adapted final interface (averaged of last 3) and the non-adaptive baseline (average of 6 baseline trials) using paired *t*-tests at each stimulated frequency (*α* = 0.05). The null hypothesis **H1**_**0**_ is: there is no significant difference in the transformations of either the interface or the humans after adaptation compared to their respective baselines.

###### Algorithm 1

Interface update during co-adaptation

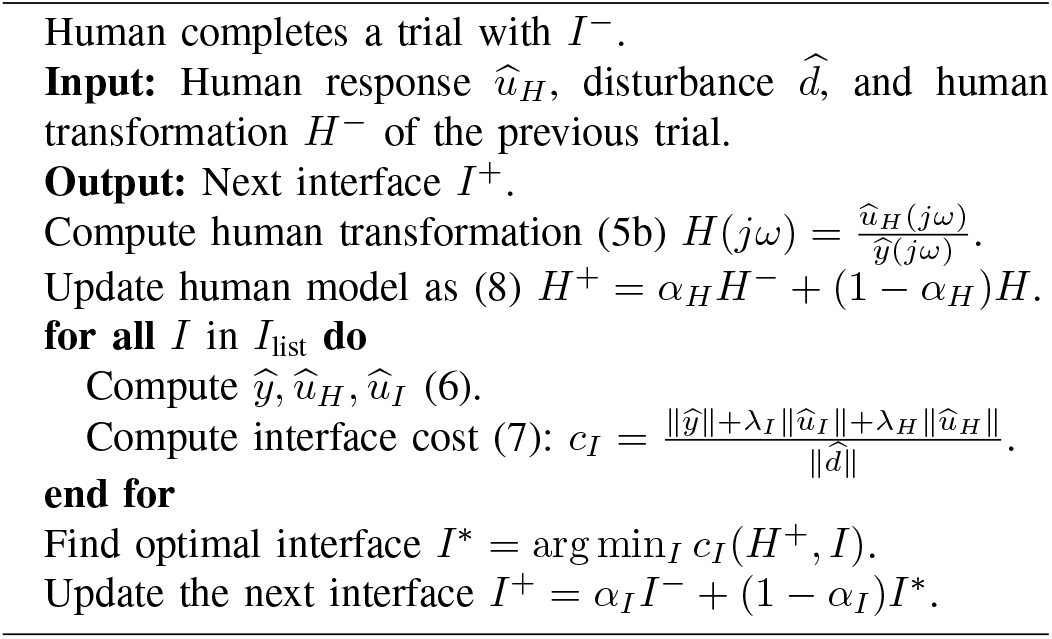

##### H2 (baselines vs. pass-through)

We investigated whether the three baselines (*I*_0,base_, *I*_1,base_, *I*_2,base_; Table I) improved from having “no interface” (pass-through with *I* = 1, *I*_pass_) in terms of the interface cost *c*_*I*_ (*H, I*) defined in (7). We first confirmed normality of the average cost of each group using Shapiro-Wilk test (*p >* 0.05), then performed one-way ANOVA to compare the groups. We used paired *t*-tests with Bonferroni correction as post-hoc tests. The null hypothesis **H2**_**0**_ is: there is no significant difference in performance between baselines and pass-through.

##### H3 (adaptive vs. non-adaptive)

Our primary outcomes of interest were the differences in performance with and without co-adaptation. Because our algorithm explicitly optimizes the interface cost, we identified the reduction of this cost as the primary indicator of performance improvement. Additionally, we assessed changes in task error, human effort, and interface effort to reflect the practical goal of maximizing task performance while minimizing burdens from both agents. We defined our performance metrics as follows:

1. **Task error**: norm of the cursor displacement 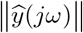 at the stimulated frequencies of a trial.
2. **Human effort**: norm of the manual device inputs 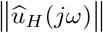 at the stimulated frequencies of a trial.
3. **Interface effort**: norm of the interface inputs 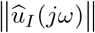 calculated in (5a) at the stimulated frequencies of a trial.
4. **Interface cost**: *c*_*I*_ (*H, I*) defined in (7) at the stimulated frequencies of a trial.
5. **Stability margin**: the stability margin of the open loop transfer function *L*(*jω*) = *H*(*jω*)*M* (*jω*)*I*(*jω*), by calculating the absolute distance between *L*(*jω*) and the critical point 1 in Nyquist plot. Higher value indicates better stability margin. We confirmed normality using Shapiro-Wilk test (*p >* 0.05), and used paired *t*-tests to compare the co-adapted final trials (average of last 3) and baseline trials (average of 6) in each performance metric, for each condition. The null hypothesis **H3**_**0**_ is: there is no significant difference in each performance metric between co-adapted finals and baselines.

We used Python 3.11 for Shapiro-Wilks tests, ANOVA test, and paired *t*-tests. Data and analyses to reproduce the results are available in a publicly-accessible repository [54].

### G. Human system identification

To characterize co-adaptation in simulation, we needed to first identify a parsimonious model of the human feedback controller in post-hoc. In a previous study, we estimated the human controller in 2nd-order machine interaction using the crossover model in [1]:

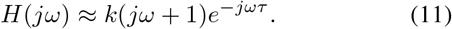

We estimated the human gain *k* and sensorimotor delay *τ* from prior experimental data in [3], which had the same disturbance-rejection task but a simpler 2nd-order machine dynamics. We reported in [55] with the average human gain and delay of *k* = 2.75 and *τ* = 0.321 seconds, respectively.

In this study, since finite-order interfaces were introduced into the system, the crossover model with a 2nd-order machine may no longer apply. However, humans do not implement arbitrarily high-order transformations [1], [43]. Therefore, we started with models incorporating the lowest possible degrees of dynamics: 1st- and 2nd-order dynamics, each with a human sensorimotor delay of *τ* = 0.321:

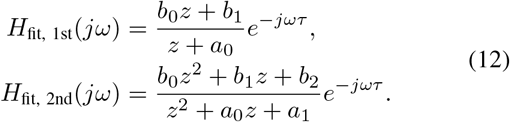

We fit the 1st- and 2nd-order models to the empirical human transformation by minimizing the difference between the modeled and empirical data. Specifically, we performed a grid search of parameters *a, b* over the range [−10, 10], and performed optimization with the L-BFGS algorithm.

### H. Co-adaptation simulation

Building upon our theoretical framework and experimental observations, we conducted a simulation to derive a computational model of the HMI co-adaptation. The objective was to identify the simulation parameters that best approximate our empirical results. Since our theory showed that there can be no Nash equilibrium when the order of the human and interface dynamics are unbounded, we constrained the simulation to low-order system dynamics for both human and interface. We evaluated the fit for both 1st- and 2nd-order human models with sensorimotor delay (Section III-G), and selected the model order that most accurately captured the empirical human transformations. Following prior work [37], [53], we modeled human adaptation as the minimization of a cost function defined as a linear combination of task performance and human effort, mirroring the structure of the interface cost:

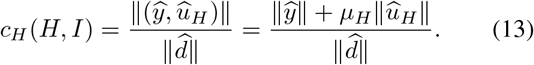

We applied an alternating optimization procedure to synthesize the optimal human *H*^∗^ and interface *I*^∗^ in simulation. Starting with the baseline *I*_base_ as the initial interface, we synthesized *H*^∗^ by minimizing human cost in (13). We imposed constraints on the human model to reflect physiological limitations: a phase constraint of ∠*H < π* because humans cannot predict future task signals, and a magnitude constraint of |*H*| ≤ 2.5 for frequencies ≥0.55 Hz to account for the limited human control bandwidth. Human parameters *b*_0_, *b*_1_, *b*_2_, *a*_0_, *a*_1_ in (12) were randomly initialized over the range [−20, 20] and the optimal human *H*^∗^ was obtained by minimizing the non-convex cost using the Sequential Least Squares Programming (SLSQP).

Subsequently, we held *H*^∗^ constant and synthesized *I*^∗^ by minimizing the interface cost *c*_*I*_ (*H, I*) using the experimental algorithm. This alternating optimization was iterated 15 times, mirroring the 15 trials performed in the empirical study.

We performed a parameter sweep of the penalty terms, *λ*_*I*_, *λ*_*H*_ in (7) and *µ*_*H*_ in (13), across the range [0.5, 10]. The optimal set of penalties was identified by minimizing the difference between the simulated and the experimental data. Model performance was evaluated using the standardized distance (*z*-score): 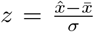, where 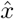 denotes the simulated value, and 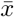 and *σ* represent the mean and standard deviation of the empirical subject data, respectively. A simulated result was considered consistent with experimental result if |*z*| *<* 1.

## IV. Experimental Results

Eleven participants completed the disturbance-rejection task in the presence of a dynamic machine in (4), over a sequence of 21 trials (3 non-adaptive baselines; 15 co-adaptations; 3 non-adaptive baselines) in each of the three conditions presented in random order: 0th-, 1st, and 2nd-order interfaces (Table I).

### A. Humans and interfaces co-adapted

We tested our first hypothesis, whether humans and interfaces co-adapted in our experiment (**H1**). We found significant differences between the magnitudes of non-adaptive baseline interfaces (dashed lines; Figure 4a) and the co-adapted final interfaces (blue lines; Figure 4a), where *p <* 0.05 at all eight stimulated frequencies. This implies that in all three conditions, after 15 trials of co-adaptation, interfaces adapted to a different interface than the baseline.

**Fig. 4.**
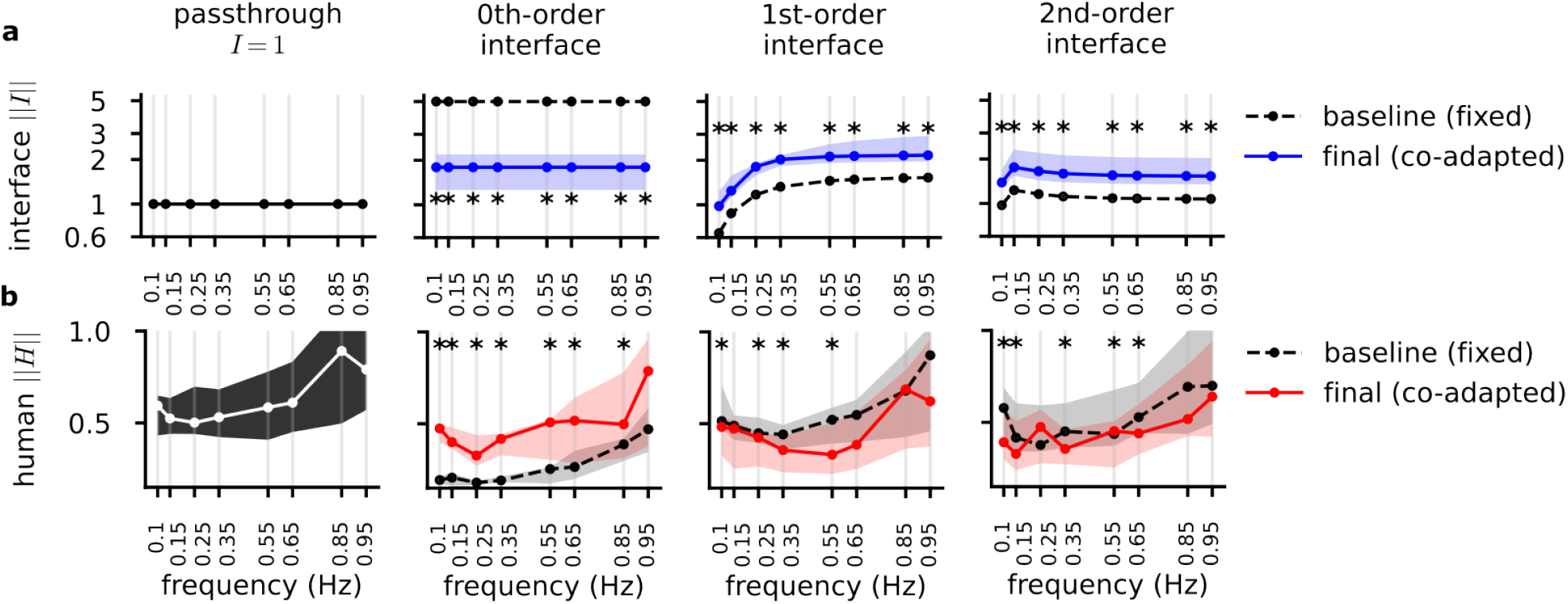
Experiment results on human-interface co-adaptation. Spectral density plots (median and interquartile, *N* = 11 participants) of **(a)** magnitudes of pass-through interface (*I* = 1), non-adaptive baseline interfaces (dashed lines), and co-adapted final interfaces (blue lines) for the 0th-, 1st-, and 2nd-order conditions at the stimulated frequencies. **(b)** Magnitudes of human transformations in pass-through trials, non-adaptive baseline trials (dashed lines), and the co-adapted final trials (red lines) for the 0th-, 1st-, and 2nd-order interfaces. Statistically significant differences between baselines and co-adapted final transformations are shown with an asterisk (^∗^ *p <* 0.05, paired *t*-test at each stimulated frequency).

We then examined human adaptation in response to these interface adaptations. We found significant differences between human transformations in baseline trials (dashed lines; Figure 4b) and co-adapted final trials (red lines; Figure 4b). These differences *p <* 0.05 were present across the majority of stimulated frequencies, particularly in the low-frequency range (*<* 0.55 Hz). Thus, we reject the null hypothesis **H1**_**0**_.

In addition, we observed a trade-off in magnitude shifts of the human and interface agents. In the 0th-order condition, interface magnitudes decreased relative to baseline while human magnitudes increased. Conversely, in the 1st- and 2nd-order conditions, we observed that interface magnitudes increased, whereas human magnitudes decreased. These results provide evidence of a trade-off between the two decision-making agents as they co-adapt in this dynamic game.

### B. High pass interfaces improved performance

We tested our second hypothesis, whether the baseline interfaces would yield a different performance than the pass-through interface (**H2**). We defined our primary performance metric as the interface cost *c*_*I*_ (*H, I*) in (7).

One-way ANOVA with post-hoc paired *t*-tests found no statistically significant difference in interface cost between the 0th-order baseline and the pass-through (*t*(10) = 0.097, corrected *p* = 1). Furthermore, participants subjectively reported in 0th-order baseline trials that the cursor was overly “sensitive”, likely due to the large input scaling of 5 (*I*_0,base_ = 5).

In contrast, interface costs for the 1st- and 2nd-order baselines were significantly lower than those of the pass-through condition (1st-order: *t*(10) = 8.76, corrected *p <* 0.001; 2nd-order: *t*(10) = 9.98, corrected *p <* 0.001). These findings demonstrate that the order of the interface dynamics influenced system performance, with higher-order (1st- and 2nd-order) high-pass interfaces yielding better performance relative to the pass-through baseline. Thus, we fail to reject the null hypothesis **H2**_**0**_ for the 0th-order condition, and reject the null hypothesis for the 1st- and 2nd-order conditions.

### C. Performance improved or maintained after co-adaptation

We evaluated the effect of co-adaptation using five performance metrics: task error 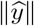 (cursor displacements), human effort 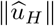 (human responses), interface effort 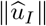 in (5a), interface cost *c*_*I*_ (*H, I*) in (7), and the stability margin of the open-loop transfer function *L* = *HMI*. We tested our third hypothesis, whether the co-adapted interfaces yielded a different performance than the non-adaptive baselines (**H3**).

In the 0th-order condition, task error (Figure 5a) was significantly reduced in the co-adapted final interface compared to the 0th-order baseline (*p <* 0.001). For the 1st- and 2nd- order interfaces, while the median task errors showed a slight decrease, these differences did not show statistical significance (1st-order: *p* = 0.14; 2nd-order: *p* = 0.65). Given that 1st- and 2nd-order baselines significantly outperformed the pass-through condition in task error, these findings suggest that the 1st- and 2nd-order baselines already yielded relatively good task performance, and that task performance was maintained after co-adaptation.

**Fig. 5.**
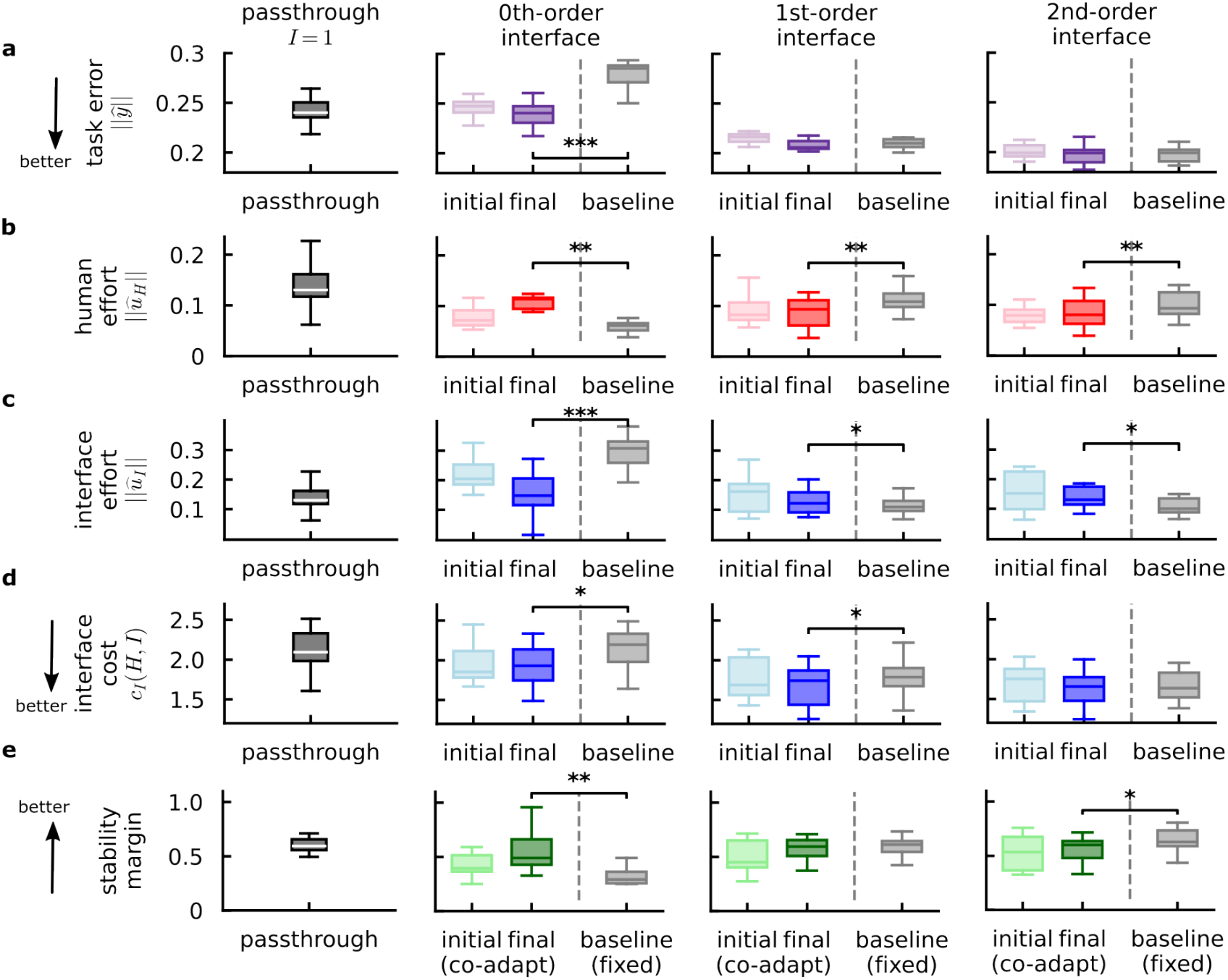
Experiment results on performance. Boxplots (0th, 25th, 50th, 75th, and 100th percentiles, *N* = 11 participants) of each performance metric for pass-through (average of 12 trials), baselines (average of 6 trials), and adaptive interfaces (average of initial 3 trials and average of final 3 trials). **(a)** task error 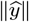 (lower values are better), **(b)** human effort 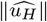, **(c)** interface effort 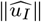, **(d)** interface cost *c*_*I*_ (*H, I*) defined in (7) (lower values are better), and **(e)** stability margin (higher values are better). Statistically significant differences are shown with horizontal lines and asterisks (^∗^ for *p <* 0.05, ^∗∗^ for *p <* 0.01, and ^∗∗∗^ for *p <* 0.001; paired *t*-test).

We observed that human effort and interface effort exhibited divergent trends across conditions (Figure 5b, c). Human effort was significantly higher in the co-adapted final interface compared to the baselines for the 0th-order condition (*p <* 0.01), whereas it was significantly reduced in both the 1st- and 2nd-order conditions (1st-order: *p <* 0.01; 2nd-order: *p <* 0.01). In contrast, interface effort was significantly lower in the co-adapted final interface compared to the baselines for the 0thorder condition (*p <* 0.001), and was significantly higher in both the 1st- and 2nd-order conditions (1st-order: *p* = 0.044; 2nd-order: *p* = 0.048).

Regarding the interface cost (Figure 5d), which was the metric explicitly minimized by our algorithm, we observed a significant reduction for the 0th- and 1st-order co-adapted interfaces compared to their respective baselines (0th-order: *p* = 0.013; 1st-order: *p* = 0.048). No statistically significant difference was observed for the 2nd-order condition (*p* = 0.1).

We evaluated robustness by comparing the stability margin of the open-loop transfer function *L* between the co-adapted final trials and the non-adaptive baselines (Figure 5e). we observed a significantly higher stability margin for the 0th- order co-adapted interface compared to the 0th-order baseline (*p <* 0.01). No significant difference was identified for the 1st-order condition (*p* = 0.29). For the 2nd-order condition, however, the co-adapted interface demonstrated a significant reduction in stability margin relative to the baseline (*p* = 0.03).

In summary, co-adaptation improved or maintained some system performance metrics, including task error and interface cost. The effort between human and interface agents shifted variably across conditions. However, co-adaptation did not necessarily enhance robustness. These findings led us to reject the null hypothesis **H3**_**0**_.

## V. Computational results

Following the system identification procedure (Section III-G), we modeled the human dynamics using both 1st- and 2nd-order system architectures with sensorimotor delay. We found that the 2nd-order model in (12) fit the empirical distribution well (Figure 6a). The two distributions (magnitudes of *H*_fit_ and *H*_exp_) exhibited Pearson correlation coefficients of at least *r >* 0.88 at every stimulated frequency (*p <* 0.001). The 1st-order model had a relatively poorer fit than the 2nd-order model, with Pearson correlation coefficients of at least *r >* 0.77 at every stimulated frequency (*p <* 0.01).

**Fig. 6.**
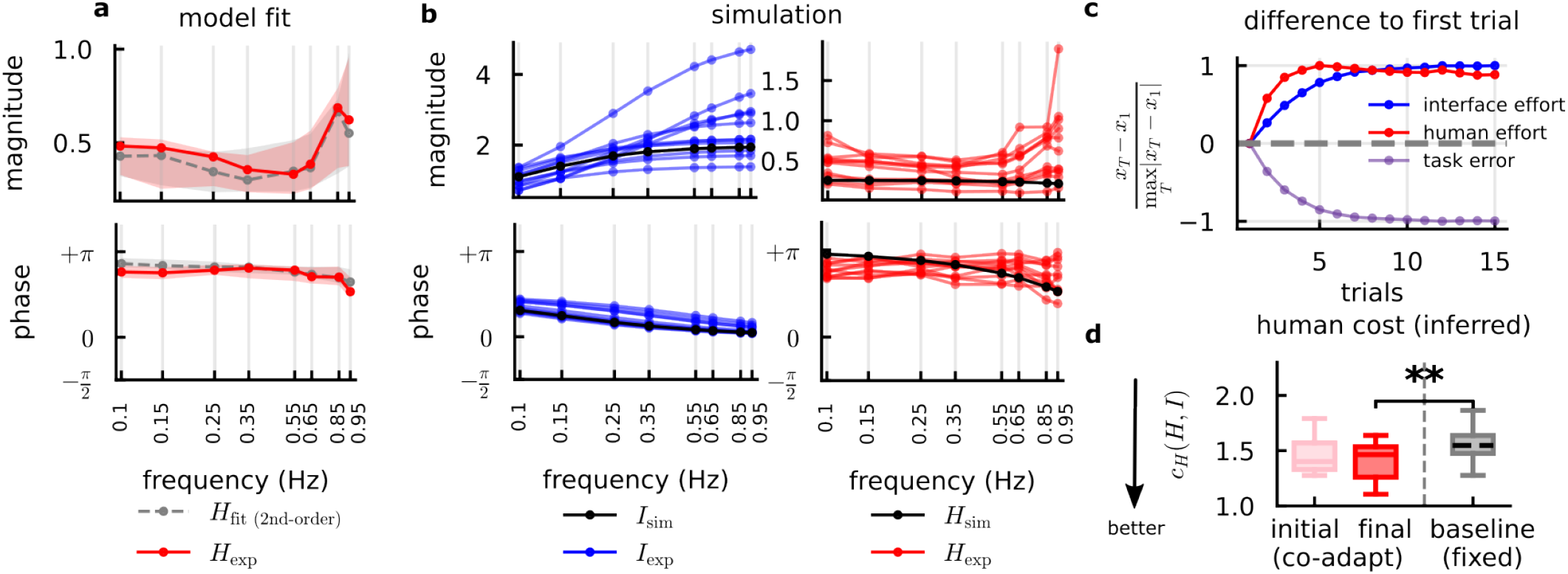
Computational results with a 1st-order interface. **(a)** Spectral density plots with distributions (median and interquartile, *N* = 11 participants) of (top) magnitudes and (bottom) phases of the 2nd-order model fit *H*_fit_ and the empirical human transformation *H*_exp_. **(b)** Spectral density plots of the (left) simulated interface *I*_sim_ and (right) simulated human *H*_sim_ overlaid on empirical data *I*_exp_, *H*_exp_ of every subject (*N* = 11 participants). Each line is the average of 3 co-adapted final trials from one of the subjects. **(c)** Difference between each trial *T* and the first trial *T* = 1 in simulation, for each element in the interface cost defined in (7): interface effort, human effort, and task error. Trial-wise metrics were plotted as deviations from the first trial, normalized by the maximum deviation, so the largest absolute change from baseline corresponds to ±1. **(d)** Boxplot (0th, 25th, 50th, 75th, and 100th percentiles, *N* = 11 participants) of the inferred human cost, with the *µ*_*H*_ value found in simulation, of 1st-order baseline and the 1st-order adaptive trials (initial and final). Statistically significant difference is shown with a horizontal line and asterisks (^∗∗^ for *p <* 0.01; paired *t*-test).

We therefore used the 2nd-order human model with sensorimotor delay to simulate a human-like agent interacting with the 1st-order adaptive interface in the experiment. In this simulation, we modeled human behavior by adopting a cost function analogous to the interface agent’s optimization strategy, defined in (13). We simulated the co-adaptation process exclusively for the 1st-order interface. This selection was based on experimental findings that only the 1st-order interface achieved significant improvements in interface cost relative to both the baseline (*p* = 0.048) and the pass-through interface (*p <* 0.001) (Figure 5d).

We conducted a parameter sweep over the penalty terms (*λ*_*I*_, *λ*_*H*_ in (7), and *µ*_*H*_ in (13)) in range [0.5, 10] and identified the optimal configuration that minimized the deviation from empirical data (*λ*_*I*_ = 1.32, *λ*_*H*_ = 5.5, *µ*_*H*_ = 2.04). The fit was validated by calculating the standardized distance (*z*-score) between the simulated and experimental magnitudes at each stimulated frequency. We found that |*z*| *<* 1 across all stimulated frequencies for the interface magnitudes (∥*I*∥, blue in Figure 6b). Similarly, for the human magnitude (∥*H* ∥, red in Figure 6b), |*z*| *<* 1 across all frequencies except 0.85 Hz. These results indicate that the simulated result consistently remained within one standard deviation of the experimental mean, thus demonstrating good correspondence between the model and empirical outcomes.

We analyzed the evolution of the cost function components 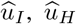, and 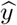, normalized to each trial’s difference to the initial trial (Figure 6c). Simulation results indicate that as interface effort gradually increased and task error decreased, human effort increased and then declined slightly, with the system reaching equilibrium after 15 trials of co-adaptation. These simulation trends demonstrated the effort and error trade-off.

With the penalty term for human effort (*µ*_*H*_ = 2.04), we computed the inferred human cost for the 1st-order interface condition in our experiment. It is important to note that these values represent model-based inferences rather than direct measurements from the experimental data. We observed that the inferred human cost significantly decreased after co-adaptation compared to the non-adaptive baseline (Figure 6d). This finding demonstrates that our proposed framework with a specific, simplified human cost function structure replicated the observed experimental outcomes.

## VI. Discussion

Machine learning algorithms are being deployed in-the-loop with people [56]. It is important to predict the outcome of these interactions to guarantee safety and performance in disparate applications including brain-machine interfaces [13], [16], [53], human-robot interaction [6], [22], [23], [33] and assistive devices [8]–[12], [15]. These interactions are increasingly viewed through the lens of game theory [4], [7], [19], [20], [32], [33], [37], [39], [53], [57], [58], where it is hypothesized that the human’s behavior arises from the solution of an optimization problem [27]–[30], [59] and the adaptive agent similarly optimizes its own cost. However, prior work makes assumptions on the ability of people to solve these optimization problems that we regard as unreasonable.

In this study, we sought to systematically test the outcome of HMI co-adaptation and compare experimental results with predictions from theory and simulations. We adopted a human-in-the-loop control paradigm wherein participants stabilized a complex (nonminimum-phase) control system through a dynamic interface as in Figure 3. This paradigm was intended to instantiate the fundamental challenges inherent in more embodied applications involving neural measurements or physical interaction with a robot or assistive device. Our testbed enables complete control over the system dynamics and external disturbances that facilitate precise comparison of experimental results with theoretical and computational predictions.

### A. Limitations of theoretical predictions

A leading hypothesis in the human motor control literature is that people solve an ℋ_2_ optimal control problem and implement a *linear-quadratic Gaussian* (LQG) controller to move their bodies [27], [28]. Prior work on co-adaptation [32]–[34] specialized this model further by considering the *full information* case where the human is assumed to have noise-free observations of the system state. The full information case has the appeal that it admits a Nash equilibrium defined in terms of static state feedback matrices [45, Sec. 6.2.2], but we argue that it is unrealistic to assume each agent has perfect knowledge of both the environment and the other agent.

Considering the more general model without full information (or full control), we find a different theoretical objection: if both the human and adaptive system solve ℋ_2_ optimization problems, there can be no Nash equilibrium for the co-adaptive system defined in terms of finite-order LQG controllers. The reason for this is a simple counting argument: since an LQG controller generally has the same order as the system it controls, and since the system one agent controls includes the dynamics of the other agent as in Figure 2, it is not possible for these system orders to “add up” to consistent quantities. Of course the Nash equilibrium is not the only candidate outcome [39]. But this thought experiment reveals an even more fundamental objection to this model for human behavior: it is simply not plausible that people are able to implement LQG controllers of arbitrary system order – they must have bounded rationality [59], [60]. These observations motivated us to conduct an experiment with adaptive systems that had fixed system orders toward the goal of understanding how people actually play this class of human-in-the-loop game.

### B. Co-adaptation shaped behavior and performance

We observed co-adaptation in our experiments: the baseline and final co-adapted human and interface transformations were different for all interface orders in Figure 4. Of course, the interface adapts by design – it repeatedly solves the optimization problem in (9). But there is no guarantee that (i) the human will adapt in response and (ii) the outcome of this interaction will differ from baseline. The fact that we observed both (i) and (ii) is particularly significant since we started with a baseline that was synthesized using prior knowledge of human behavior [55], i.e., our baseline was not naïve.

Our experimental results demonstrate potential advantages – and drawbacks – of deploying adaptive interfaces in-the-loop with humans and dynamic machines. Since our interface sought to minimize the cost in (7), it seems natural to expect this cost to decrease in the experiment. However, this decrease is not guaranteed due to the presence of human adaptation, as co-adaptive interactions may converge to a constellation of equilibrium outcomes [39]. We observed significant decreases in this cost for the 0th- and 1st-order interfaces but not for the 2nd-order interface (Figure 5d), confirming performance gains are not guaranteed.

A similar observation holds for the stability margin of the closed-loop system: solving the interface’s ℋ_2_ optimization problem can yield arbitrarily small margins in theory [44]. Experimentally, we found a significant increase, decrease, or no change in the stability margin across the different interface orders (Figure 5e). Future work may seek to prioritize robustness using ℋ_∞_ controller synthesis [31], [41], [47], [61]. A consistent finding across the three interface conditions was that the human and interface efforts traded off, as an increase in one was accompanied by a decrease in the other (Figure 5b,c). This outcome comports with other theoretical and experimental findings [37], [53] and supports the hypothesis that the human and interface play a *game* [21], [39] whose outcome represents a tradeoff between competing objectives.

### C. A computational model for human adaptation

We found that the transformations our human subjects implemented in-the-loop with a 1st-order interface could be well-approximated as a second-order system with sensorimotor delay (Figure 6a). Importantly, the dimension of the state for the human’s transformation *H* in this case (2) is less than the combined dimension of the machine *M* and interface *I* (2 + 1 = 3), undermining the hypothesis that the human implements the LQG controller whose state dimension would match that of the combined machine-interface system *M/I*. This finding is consistent with prior work and supports the hypothesis that humans do not implement transformations of arbitrarily-high order [1], [43].

We found that modeling the human as optimizing a cost function consisting of a linear combination of error and effort – inspired by Emken *et al*. [30] – can produce similar outcomes to what we observed in our experiment (Figure 6b) and yield an error-effort tradeoff (Figure 6c). We regard this result as provocative but far from definitive. In particular, we do not believe our human subjects literally optimized a cost function with the precise parameterization in (13), as human motor control can be influenced by cognitive load, fatigue, injury, or distractions [62]–[64]. All models are wrong, but some are useful [65]; interrogating the utility of our proposed model is an important goal for future work.

### D. Principles for interface design

Motivated by the observation that human sensorimotor delay constrains performance in trajectory-tracking tasks [61], we employed high-pass interfaces in our experiments. Indeed, the human brain attempts to compensate for delay internally through feedback projections, but this compensation is imperfect and is subject to speed-accuracy trade-offs [66]. Adding a high-pass interface can supplement this phase-lead mechanism in the human’s internal delay compensation, potentially improving performance of the human-in-the-loop system. Figure 5a,d demonstrates that the high-pass baselines (1st- and 2nd-orders) and co-adapted final interfaces decreased task error and interface cost compared to the pass-through.

If human behavior in co-adaptive interactions arises from optimization as corroborated by our simulation results, it opens the possibility of systematically shaping interaction outcomes by modifying the interface cost function. A proof-of-concept of this idea was demonstrated in [53], where it was shown that increasing a penalty term in the interface cost decreased interface effort and increased human effort. This observation has broad implications for the design of both assistive devices, which seek to minimize user effort, and rehabilitation devices, which aim to promote active patient engagement [67], [68]. Game-theoretic methods that explicitly consider the strategic interests of both decision-making agents may enable designers to systematically select from a variety of outcomes [39].

## VII. Conclusion

Our work highlights limitations in prior theory, experiment, and simulation work on co-adaptation between humans and intelligent interfaces interacting in closed loop with dynamic machines in real-world conditions. We demonstrate that a Nash equilibrium does not exist under standard assumptions of optimal adaptation, and explore in experiment and simulation how a human and interface with *bounded rationality* [59] adapt to improve performance in a disturbance-rejection task. Understanding how to analyze and synthesize co-adaptive systems, where both human and algorithmic agents adapt to control dynamic systems like vehicles, robots, and prostheses is critically important in current and emerging applications including driver assistance, rehabilitation robotics, and neuroprosthetics. We contribute new theory, experiment, and simulation results that will inform creation of these systems for real-world deployment.

## Acknowledgments

We thank the participants in our experiment. This work was funded by a American Society for Engineering Education National Defense Science and Engineering Graduate Fellowship (NDSEG) to MMM and National Science Foundation (NSF) awards #2045014,2124608 to SAB.

